# Pattern formation by bacteria-phage interactions

**DOI:** 10.1101/2023.09.19.558479

**Authors:** Alejandro Martínez-Calvo, Ned S. Wingreen, Sujit S. Datta

**Affiliations:** Princeton Center for Theoretical Science, Princeton University, Princeton, NJ 08544, USA; Department of Chemical and Biological Engineering, Princeton University, Princeton, NJ 08544, USA; Lewis-Sigler Institute for Integrative Genomics, Princeton University, Princeton, NJ 08544, USA; Department of Molecular Biology, Princeton University, Princeton, NJ 08544, USA

## Abstract

The interactions between bacteria and phages—viruses that infect bacteria—play critical roles in agriculture, ecology, and medicine; however, how these interactions influence the spatial organization of both bacteria and phages remain largely unexplored. Here, we address this gap in knowledge by developing a theoretical model of motile, proliferating bacteria that aggregate via motility-induced phase separation (MIPS) and encounter phage that infect and lyse the cells. We find that the non-reciprocal predator-prey interactions between phage and bacteria strongly alter spatial organization, in some cases giving rise to a rich array of finite-scale stationary and dynamic patterns in which bacteria and phage coexist. We establish principles describing the onset and characteristics of these diverse behaviors, thereby helping to provide a biophysical basis for understanding pattern formation in bacteria-phage systems, as well as in a broader range of active and living systems with similar predator-prey or other non-reciprocal interactions.

---

Bacteriophages (“phages”) are viruses that infect bacteria. This predator-prey relationship regulates how bacterial communities form and function in diverse environments, ranging from the ground beneath our feet to the tissues and organs in our bodies [1–6]. Thus, it is important to understand how interactions with phages shape bacterial populations. Indeed, these interactions are of fundamental interest in biology, ecology, and physics, and have critical implications for agriculture, ecology, and medicine [4–19].

Studies using continuously-mixed liquid cultures have yielded crucial insights into bacteria-phage interactions [5, 20–27]. However, many bacteria inhabit more quiescent environments where the cells can collectively self-organize into spatially structured populations. Despite the pivotal influence of bacterial spatial organization on diverse biological functions, how such organization is influenced by, and in turn influences, interactions with phages remains poorly understood.

Here, we address this gap in knowledge by developing a minimal theoretical model that incorporates essential biophysical features of bacterial spatial organization and interactions with phage. In particular, we describe the bacterial population as a collection of actively-moving and proliferating particles that tend to undergo motility-induced phase separation (MIPS). In this collective self-trapping process, the bacterial cells slow down where they form aggregates, and aggregate where they are slower. This feedback loop makes the bacterial population unstable to density fluctuations, leading it to spatially organize into dense aggregates in a manner reminiscent of phase separation via spinodal decomposition [28–30]. In addition to being a well-studied canonical model of active matter, MIPS-like behavior has been reported in many different bacterial systems [31–39], making it a useful model of bacterial spatial organization. We introduce phage into this model as additional passive particles that can diffuse, infect bacteria, proliferate inside them, and kill cells via lysis, generating a burst of new phage progeny (Figure **1**a). These bacteria-phage interactions are therefore inherently *non-reciprocal*: an asymmetrical relationship exists between the two entities, in which bacteria (the prey) are beneficial to phage, but phage (the predators) are detrimental to bacteria. Since other forms of non-reciprocal interactions can lead to pattern formation, dynamical spatial structures, and phase transitions [40–42], e.g., flocking phenomena [43–46] and synchronization of living and synthetic systems [47–50], we asked: how might interactions with phages impact bacterial aggregation?

**Fig. 1.**
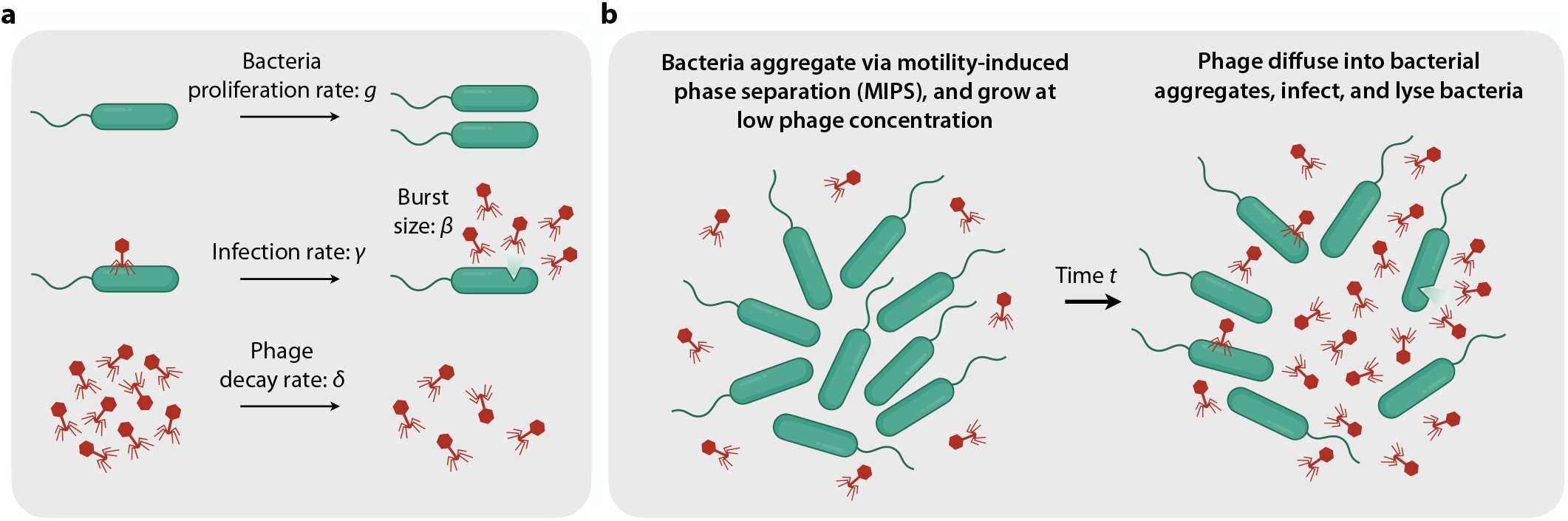
Overview of our model of bacteria-phage interactions in a spatially-extended environment. **a**. (Top) Bacteria proliferate at a rate *g*. (Middle) Phage infect bacteria at a rate *γ* and induce lysis, generating a burst *β* of phage progeny. (Bottom) Phage decay at a rate *δ*. **b**. Schematic of bacteria aggregating via MIPS at sufficiently low phage density, upon which phage can diffuse in, infect and lyse the cells, and thereby release more phage.

Using linear stability analysis and full numerical simulations of our model, we find that the dynamics of bacterial MIPS are dramatically altered by interactions with phage. At very low or very high cell densities, the bacteria remain uniformly dispersed. By contrast, over a wide range of intermediate densities, bacteria-phage interactions engender a wealth of finite-sized patterns in the bacterial population that are fundamentally distinct from the spinodal decomposition observed in conventional MIPS. In some cases, these are stationary patterns of dense bacterial aggregates, with geometries ranging from hexagonal lattices of round aggregates to disordered stripes, that coexist with phage. In others, the patterns are highly dynamic, ranging from ordered round aggregate that oscillate asynchronously to migrating threads to large domains that oscillate quasi-independently of each other.

Our analysis establishes quantitative principles describing the onset of these diverse behaviors, and elucidates the underlying cause: they arise due to the competition between MIPS, which drives bacteria to aggregate into dense phases, and the non-reciprocal predator-prey interactions caused by phage, which cause dense aggregates of bacteria to disintegrate. Furthermore, comparison to a related, but distinct, model in which bacteria are killed by a self-secreted toxin instead of by phage shows that the auto-catalytic proliferation mechanism specific to phage is essential to the advent of these dynamic patterns. Finally, we demonstrate another consequence of the nonlinear interactions between bacteria and phage: once established, the emergent bacterial patterns persist far beyond the critical phase boundaries predicted by linear stability analysis, and exhibit excitable behavior in these regions. Thus, not only does our illustrative model uncover the rich pattern formation that can arise from bacteria-phage interactions, but it also demonstrates that such interactions give rise to two hallmarks, hysteresis and excitability, of many other complex and living systems. Taken together, these results elucidate a simple biophysical mechanism that can give rise to spatial organization in bacteria-phage systems— complementing other interesting biological mechanisms such as phage hitchhiking [7] and coevolution [10]. Moreover, this model could help provide a foundation for explaining pattern formation in a broader range of active and living systems whose constituents also exhibit predator-prey relations or other non-reciprocal interactions [40, 42, 51–55].

## Model for bacteria and phage

We consider a system of motile bacteria and lytic phage in two spatial dimensions (2D). The number densities of bacteria and phage are denoted by *b*(***r***, *t*) and *p*(***r***, *t*), respectively, where ***r*** is position and *t* is time. The conservation equations for bacteria and phage read:

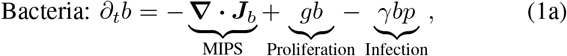

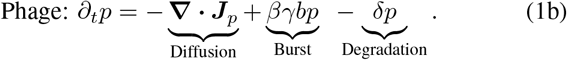

The first terms on the right in Equations (1a) and (1b) represent bacterial motility and phage diffusion, where ***J***_*b*_ and ***J***_*p*_ are fluxes of bacteria and phage, respectively. The second term in Eq. (1a) describes the proliferation of bacterial cells at a constant rate *g* reflecting nutrient-replete conditions. The third term in Eq. (1a) models bacterial infection and thus lysis and death caused by phage, where *γ* is the infection rate. For simplicity, we assume that bacterial lysis is instantaneous following infection. Hence, the second term in Eq. (1b) describes the burst of *β* + 1 phage particles arising from a lysed bacterial cell; the effective burst size is *β* since the original infecting phage is lost. Finally, the last term in Eq. (1b) reflects degradation of phage at a rate *δ*. Taken together, these last four terms describing the proliferation and removal of phage and bacteria correspond to the nonreciprocal Lotka-Volterra predator-prey model for two species [56, 57], with phage as the “predator” and bacteria as the “prey”.

The diffusive fluxes of bacteria and phage are given by:

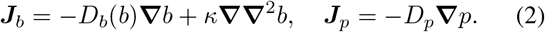

In this description, the phage diffuse uniformly across space with a diffusion coefficient *D*_*p*_. By contrast, the diffusive flux of bacteria is given by an established continuum model of MIPS [28, 30, 38] where *D*_*b*_(*b*) is an effective active diffusion coefficient reflecting undirected cellular motility, and the characteristic length scale *κ* determines the width of the interface between dense bacterial aggregates and dilute regions in MIPS.

In particular, treating the bacterial cells as run-and-tumble particles with spatially varying average speed *v*(***r***, *t*) and uniform tumbling time *τ*, the first contribution to the bacterial diffusive flux is given by ***J***_*b*,MIPS_ = −*D*_*b*_(*b*) **∇** *b* = −*v*^2^*τ***∇** *b/d* − *bvτ v/d*, where *d* is the spatial dimension, and we have assumed that the density is slowly varying [28, 58]. The first term represents standard Fickian diffusion, while the second term reflects the tendency of the cells to drift toward regions of low speed with a drift velocity *vτ***∇***v/d*. For simplicity, we assume that the spatial dependence of *v* arises only from the spatial dependence of the bacterial density *b*. Hence, ***J***_*b*,MIPS_ = −*D*(*b*) **∇***b* − *bD*^*′*^(*b*) **∇** *b/*2, i.e., *D*_*b*_(*b*) = *D*(*b*) + *bD*^*′*^(*b*)*/*2, where *D*(*b*) = *v*(*b*)^2^*τ/d* is a density-dependent diffusion coefficient, and the prime denotes a derivative with respect to *b*. In the absence of phage, such a diffusive flux gives rise to MIPS when *v*^*′*^(*b*_0_)*/v*(*b*_0_) *<* −1*/b*_0_, or equivalently *D*_*b*_(*b*_0_) *<* 0, where *b*_0_ is the spatially uniform density. As a concrete example, to obtain the numerical results presented below, we follow previous work and consider an exponential decay of the average bacterial speed with respect to density, i.e., *v* = *v*_0_ exp(−*αb/*2), where *v*_0_ is the average speed at vanishingly small density [30, 38, 59, 60]. This distribution results in an effective diffusivity *D*_*b*_(*b*) = *D*_*b*,0_ exp(−*αb*)(1 −*αb/*2), where *D*_*b*,0_ is the maximal bacteria diffusivity at vanishingly small density. However, the theoretical analysis presented below is more general, and not restricted to this choice.

## Dimensionless parameters governing our system

Before investigating how interactions with phage influence bacterial spatial organization in this model, we first reduce the number of parameters via non-dimensionalization. To this end, we choose *t*_c_ = *g*^−1^, *ℓ*_c_ = (*κ/g*)^1*/*4^, *b*_c_ = *α* ^−1^, and *p*_c_ = *g/γ* as characteristic time, length, bacterial density, and phage density scales, respectively. Non-dimensionalizing Eqs. (1a)–(1b) then yields the dimensionless Eqs. (S1a)–(S1b), revealing four key dimensionless parameters:

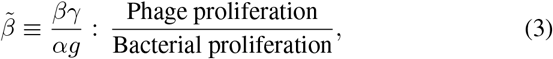

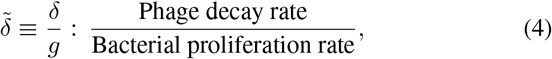

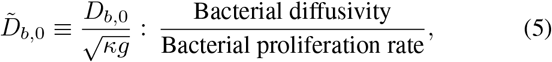

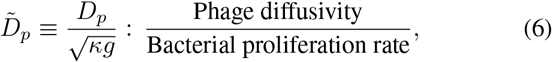

where tildes (^*∼*^) indicate dimensionless quantities. A uniform, albeit potentially unstable, solution of Eqs. (1a)– (1b) in which bacteria and phage coexist is given by (*b*_0_, *p*_0_) = (*δ/*(*βγ*), *g/γ*). In dimensionless form, this solution is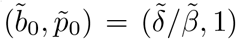, and for our subsequent analysis, we linearize the non-dimensionalized Eqs. (S1) around this solution.

## Linear stability analysis

At low phage densities, we still expect bacteria to be able to spatially organize into dense aggregates via MIPS and proliferate; however, the phage can then diffuse into these aggregates, infect and proliferate in the cells, and lyse them, releasing more phage—as depicted in Fig. **1**b. What are the possible outcomes for such bacterial aggregates and what are the consequences for the large-scale spatial organization of bacteria and phage over time?

To address this question, we analyze the linear stability of Eqs. (S1) by considering small-amplitude perturbations to the bacterial and phage densities of amplitude 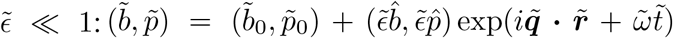, where 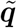 is the wave vector of a given perturbation and 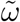 is its growth rate. Thus, the perturbation will grow if 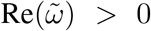, or alternatively decay if 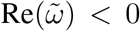. If the growth rate has an imaginary component, 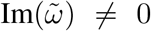, perturbations oscillate while growing or decaying over time. Introducing this expression into Eqs. (S1) yields the following dispersion relation:

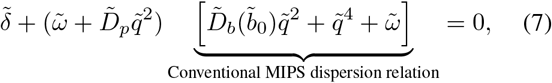

where the spatial wavenumber 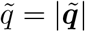. In the absence of both bacterial and phage proliferation, Eq. (7) reduces to the conventional MIPS dispersion relation, 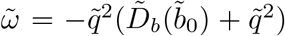. In this case, the bacterial population remains uniform below a critical density where the effective diffusivity is positive, 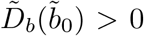, but phase separates via spinodal decomposition and coarsens over time above this density where 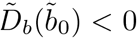. Above the critical bacterial density 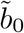, only wavenumbers 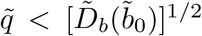have 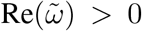 and give rise to conventional MIPS [30], as expected.

How would interactions with phage influence the spatial organization of bacteria in the case of a constant bacterial diffusivity 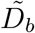that does *not* depend on cell density—that is, in a population that does not undergo MIPS? Despite superficial similarity to a Turing system [61]—with bacteria as “activator” and phage as “inhibitor”—the bacterial population remains stable, since 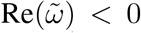for all 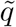, However, modes with 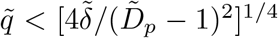, where 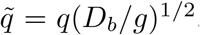, have 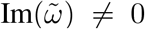, indicating that these perturbations decay over time by oscillating toward the uniform solution. This observation hints that, while in the non-MIPS case patterns do not form in the bacterial population, in the unstable case of MIPS, interactions with phage may give rise to dynamic patterns of both bacteria and phage.

## Bacteria-phage interactions can lead to the formation of stationary or dynamic patterns

To test this hypothesis, we obtain the linear stability conditions of Eq. (7). As detailed below, we first find the critical condition at which the system transitions from being stable, i.e., 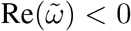for all modes 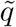, to unstable, i.e., 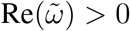for some modes, leading to pattern formation. Then, we obtain the conditions under which unstable modes also oscillate over time, i.e. 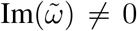. Finally, we compute the condition that determines the stability of long wavelength modes, 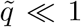. If these long-wavelength modes are unstable, dense aggregates will coalesce and coarsen over time during the linear stages when perturbations are small. Conversely, if the long-wavelength modes are stable, coarsening is prevented from progressing even within the linear regime. Nonetheless, linear stability cannot predict the full nonlinear behavior and ultimate fate of the system—therefore, to explore these fates, we also perform full time-dependent numerical simulations of Eqs. (S1).

We find that the conventional stability condition for bacterial MIPS, 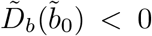, does not hold in the presence of phage. Instead, from Eq. (7) we find two distinct regions in which the uniform solution is unstable, leading to pattern for mation. First, in the region where

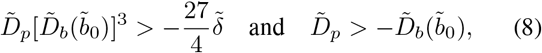

Re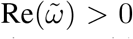for a range of 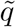with 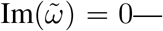indicating that the unstable modes are stationary. Moreover, in this region, 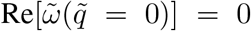 and 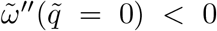, which implies that long wavelength perturbations are suppressed and will not grow; thus, we expect coarsening to be arrested in this region. Second, in the region where

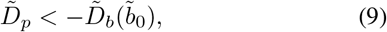

a range of unstable modes with 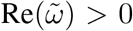exhibit a nonzero imaginary part 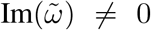, implying they are oscillatory. Since 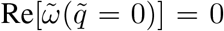and 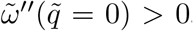, we expect the oscillating patterns that arise in this region to coarsen in the linear regime. These two linearly-unstable regions are shown by the blue and orange regions in Fig. **2**a, respectively, which characterizes the stability of the system as a function of dimensionless phage decay rate 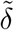and diffusion coefficient 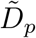.

**Fig. 2.**
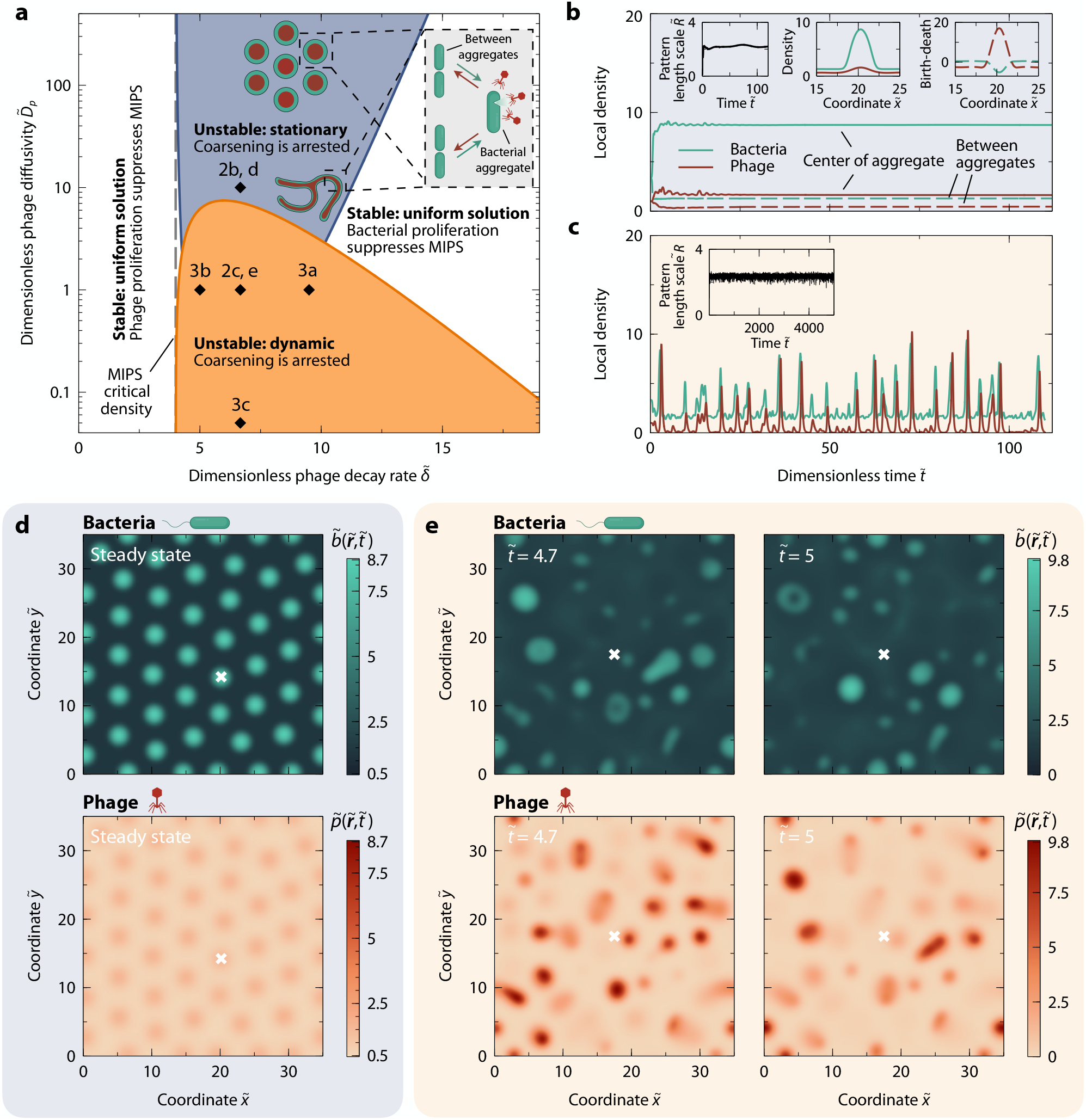
Bacteria-phage interactions can give rise to stationary or dynamic patterns. **a**. Linear stability phase diagram as a function of the dimensionless phage diffusivity 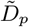and phage decay rate 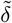. We use the bacterial density-dependent diffusivity evaluated at the uniform solution, 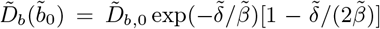, with 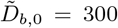, and a dimensionless phage:bacteria proliferation ratio 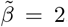we employ these physically reasonable parameter values of 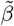 and 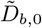 for all numerical simulations (*Supplementary Information*). The shaded blue and orange show regions where the system is unstable, with patterns that are, respectively, stationary or dynamic, and in both cases exhibit arrested coarsening. The left and right are regions where phage or bacterial proliferation, respectively, suppresses phase separation. Diamonds indicate simulations shown in panels b-e and Fig. **3. b**. Local bacteria and phage densities as a function of dimensionless time 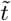for the linearly-unstable stationary case indicated in panel a 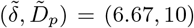. Solid and dashed curves indicate the density of bacteria and phage at the center of an aggregate and between aggregates, respectively. Insets: left, characteristic pattern length scale 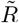 as a function of time; middle and right, bacterial and phage densities and net birth-minus-death rates across a stationary round aggregate. **c**. Same as in panel b, but for the linearly-unstable, dynamic case indicated in panel a 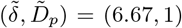. **d**. Two-dimensional plot of the bacterial and phage densities 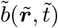 and 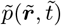, respectively, at steady state for the stationary case shown in panel b. The symbol marks where the densities in panel b are measured. **e**. Same as in panel d, but for the dynamic case shown in panel c, at times 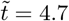and 5.

These results make sense intuitively. Phase separation via MIPS consists of a positive feedback loop as bacteria tend to accumulate where they slow down and slow down where they have accumulated, resulting in non-oscillatory aggregates that coarsen over time. However, the presence of phage introduces a negative feedback loop: aggregation and proliferation create dense regions of bacteria, favoring rapid proliferation of phage which reduces the density of bacteria. The combination of these mechanisms can, in principle, give rise to either stationary or dynamic patterns.

What happens as the instabilities grow into the nonlinear regime? To obtain the full nonlinear spatio-temporal patterns in the different linearly-unstable regimes of Fig. **2**a, we perform numerical simulations of Eqs. (S1) (*Supplementary Information*). As we describe next, we find both stationary and dynamic patterns, but in all cases, coarsening is arrested—in contrast to conventional MIPS.

## Stationary patterns emerge from the interplay of bacterial MIPS and lysis by rapidly-diffusing phage

Stationary patterns of bacteria and phage always emerge in the upper linearly-unstable region of the phase diagram in Fig. **2**a, i.e., when phage diffusivity is sufficiently high. Far inside this region, the patterns consist of a hexagonal lattice of stationary round aggregates in which bacteria and phage coexist at high density, as shown in Figs. **2**b and d (Supplementary Movies 1 and 2). The mean aggregate size 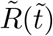, computed from Eq. (S15), plateaus at a constant value, shown in the left inset to Fig. **2**b, as expected.

What biophysical mechanism gives rise to these steady-state patterns? In the absence of phage and of bacterial proliferation, the bacterial aggregates formed by MIPS coarsen via Ostwald ripening, with an average aggregate size that grows as 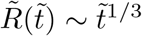 [30, 62, 63]. The non-reciprocal predator-prey interactions caused by phage modify this paradigm. In particular, after bacterial aggregates form via MIPS, phage already present at those locations or that rapidly diffuse into the aggregates kill the bacteria and proliferate. This process suppresses further coarsening of the bacterial aggregates. However, it does not completely annihilate the aggregates; there is a positive net birth rate of bacteria in the space between aggregates, shown by the data spanning 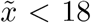and 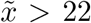in the right inset to Fig. **2**b. Thus, at steady state the killing of bacteria in aggregates is compensated by a MIPS flux of bacteria from the space between aggregates, as schematized in the inset to Fig. **2**a, along with fast diffusion of phage out of the aggregates after they proliferate.

Interestingly, closer to the critical boundaries of this region of the phase diagram in Fig. **2**a, more intricate stationary patterns emerge. For example, near the righthand boundary where phage decay rapidly (large 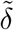), we observe the disordered bacteria-phage stripes shown in Fig. **S3**—reminiscent of the patterns that have been reported to emerge in phage-free models of MIPS that incorporate logistic growth of bacteria [38].

## Dynamic patterns emerge from the delay between bacterial MIPS and lysis by slowly-diffusing phage

Non-stationary patterns always emerge in the lower linearly-unstable region of the phase diagram in Fig. **2**a, i.e., when phage diffusivity is sufficiently low. A striking variety of dynamic patterns emerge in the different parts of this region as perturbations grow into the nonlinear regime. One example is a disordered array of round aggregates each of which forms and disappears, as shown in Figs. **2**c and e (Supplementary Movies 3 and 4); intriguingly, these dynamics are chaotic, showing no spatio-temporal correlations and exhibiting a positive maximum Lyapunov exponent (*Supplementary Information*).

We again ask: What biophysical mechanism gives rise to these patterns? As in the stationary case, bacteria aggregate via MIPS and phage infect, proliferate in, and lyse the cells in the MIPS aggregates. However, in this region of the phase diagram in Fig. **2**a, the low diffusivity of phage means that the phage density in bacterial aggregates progressively increases to the point that aggregates are split up or entirely destroyed. Subsequent bacterial proliferation then drives the formation of other bacterial aggregates later on, which are then infected by phage again—leading to ongoing cycles of bacterial aggregation followed by phage disruption, with a time delay between the two, as shown by the traces in Fig. **2**c. Similar mechanisms leading to travelling waves and migrating round aggregates have been reported previously in the context of binary mixtures with other forms of non-reciprocal interactions [51, 52, 64, 65].

Although linear stability analysis suggests that these dynamical patterns can coarsen over time as long wavelength perturbations are linearly-unstable, nonlinearities prevent the system from coarsening. For example, in the simulation shown in Figs. **2**c and e (Supplementary Movies 3 and 4), the dynamic round aggregates have a characteristic size that chaotically fluctuates around a constant value (inset to Fig. **2**c). Indeed, throughout the linearly-unstable oscillatory region of the phase diagram in Fig. **2**, we find that all patterns have a bounded characteristic length scale. Notably, this length scale is comparable to that of the mean aggregate size in the stationary patterns, suggesting that the mechanisms responsible for arrested coarsening are related in both cases.

A rich variety of dynamic patterns emerges throughout this region of the phase diagram in Fig. **2**a. For instance, when phage decay much faster than bacteria can aggregate and proliferate (large 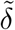), aggregates become ordered, oscillating quasi-independently in density and size, as shown in Fig. **3**a (Supplementary Movies 5 and 6). Conversely, when phage are more stable (small 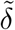), the system breaks up into finite-sized domains each of which oscillates between a uniform state in which bacterial density is low and a striped pattern, as shown in Fig. **3**b (Supplementary Movies 7 and 8). Finally, when phage diffusivity is reduced further (small 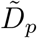), bacterial aggregates become motile, with phage pursuing the aggregates, as shown in Fig. **3**c (Supplementary Movies 9 and 10). As time progresses, these aggregates become more elongated and thread-like, also reaching a finite size.

**Fig. 3.**
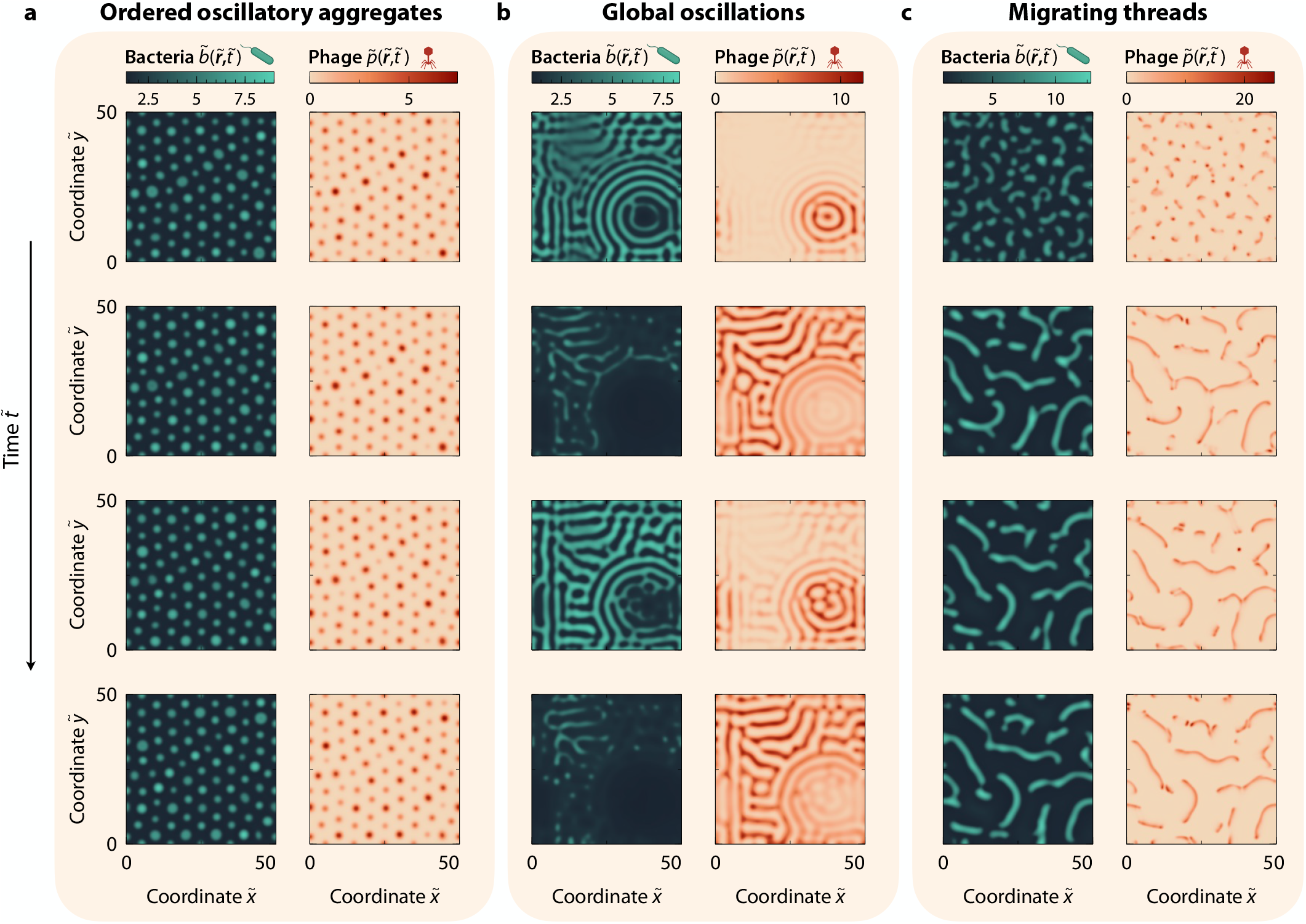
Bacteria-phage interactions give rise to a diverse array of dynamic patterns. Snapshots show bacteria and phage densities as a function of time. **a**. Ordered oscillatory round aggregates emerge when the phage lifetime is short compared to bacterial doubling time 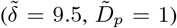. **b**. Same as in panel a but for a longer phage lifetime 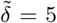in this case, large domains of striped patterns emerge and oscillate in amplitude. **c**, Same as in panel a, but for lower phage diffusivity 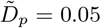and 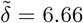 in this case, aggregates become motile and ultimately form migrating strands.

## Persistence of patterns outside linearly-stable regions

Do the observed stationary and dynamic patterns, once established, persist outside the linearly-unstable regions of the phase diagram shown in Fig. **2**a? To address this question, we conduct simulations starting just inside the rightmost phase boundaries. Once a pattern is obtained, we abruptly increase the phage decay rate 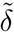, causing the system to cross the boundary into the linearly-stable region (Fig. **4**a). Beginning with a stationary case, we find that the pattern initially established indeed persists (Fig. **4**b and c)—exhibiting a similar morphology, but with a higher bacterial density within the bacterial aggregates. This makes intuitive sense, since phage proliferation slows down with increasing 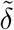. We repeat this process of increasing 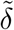 in steps, and find that stationary patterns with increasing bacterial density and thicker stripes persist up to a considerable distance into the linearly-stable region (Fig. **4**b-e). Thus, this system exhibits hysteresis, a characteristic feature of many other complex and living systems.

**Fig. 4.**
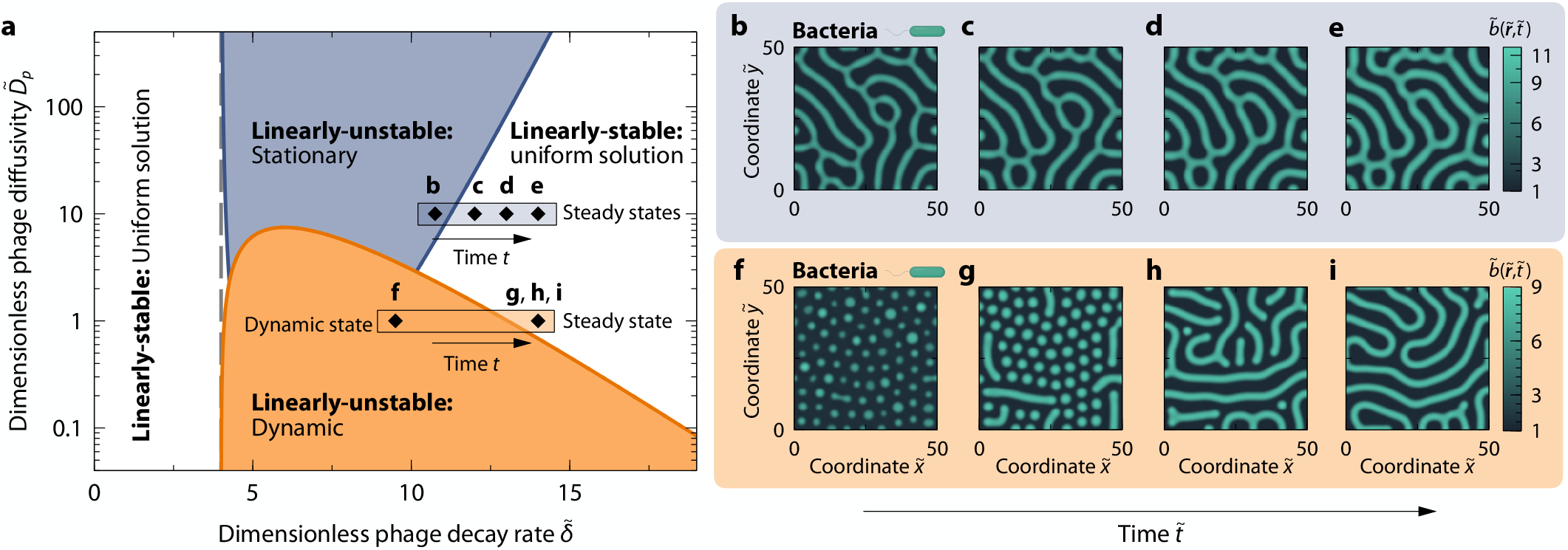
Patterns persist outside linearly-unstable regions. **a**. Same linear stability diagram as in Fig. **2**a, but showing different paths of varying 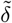 taken to investigate hysteresis, as shown by the simulations in panels b–e and f–i. After a pattern is established in a linearly-unstable region, 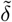 is increased suddenly and is kept constant until a new state is reached. **b**-**e**. Bacterial density 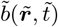 for the different values of 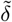 corresponding to the symbols in panel a with 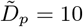, showing that stationary patterns persist outside the linearly-unstable region. **f**-**i**. Same as in panels b–e but for 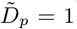, such that the system starts in the linearly-unstable dynamic region. The system transitions from a dynamic pattern of oscillatory round aggregates, as shown in Fig. **3**a, to stationary disordered stripes in the linearly-stable region. Panels g and h show intermediate states before the stationary striped pattern of panel i is reached.

We then apply the same numerical protocol starting instead with a dynamic pattern. Surprisingly, in stark contrast to the stationary case, we find that the system transitions from a lattice of oscillatory round aggregates, as in Fig. **3**a, to a stationary labyrinthine pattern of stripes (Fig. **4**f-i). In the same linearly-stable regions, we also find that the system is excitable in response to finite perturbations (Supplementary Movies 11-14)—another hallmark of many complex and living systems. This persistence of patterns outside the region of linear instability is analogous to phase separation occurring outside of the region of spontaneous spinodal decomposition in other systems [41, 66–71]. Finding the corresponding “binodal” boundaries beyond which patterned solutions no longer exist for this intrinsically nonequilibrium system will be a useful topic for future exploration.

## The auto-catalytic nature of phage proliferation is necessary for dynamic pattern formation

Bacteria can be killed by many other agents beyond phages, such as self-secreted toxins [72, 73] and waste products [74]. These other forms of killing bear some similarities to killing by phage, in particular increasing with bacterial density, but with one notable difference: phage proliferate in an auto-catalytic manner. How important is this feature of phage to the patterns that emerge in our model? To address this question, we use a similar approach to examine an alternate version of our model: instead of being killed by phage with a resulting burst of phage progeny, bacteria secrete a self-harming toxin at rate *ν* (*Supplementary Information*). Hence, the only difference between this model and our main bacteria-phage model lies in the phage proliferation term, which is instead replaced by a toxin production term proportional to the bacterial density only, i.e., *νb*. In this alternate case, dynamic patterns do not arise in the equivalent 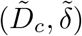 parameter space (Fig. **S1**), where 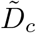 is now the dimensionless toxin diffusivity. Instead, we only observe stationary patterns with structures similar to those found for the bacteria-phage model. Thus, the auto-catalytic nature of phage proliferation is necessary for the formation of the dynamic patterns shown in Figs. **2**e and **3**.

## Discussion

In nature, bacteria and phages coexist in spatially-extended environments, such as in hosts and terrestrial and marine settings. Thus, phage predation can reshape the spatial organization of a bacterial population, which in turn alters subsequent phage infection—giving rise to an intricate array of coupled nonreciprocal predator-prey interactions across time and space whose implications remain largely unexplored. In this paper, we developed a theoretically-tractable model that describes how bacteria-phage interactions strongly impact bacterial aggregation, using the canonical model of MIPS as an illustrative example. Through a combination of linear stability analysis and numerical simulations, we showed that fascinating spatiotemporal patterns emerge from these coupled processes, and established quantitative rules describing their onset and characteristics. Our work thus sheds light on how spatial organization can arise from simple biophysical interactions between bacteria and phage. It thus complements other studies focusing on other interesting biological mechanisms, such as phage hitchhiking [7] and coevolution between bacteria and phages [7, 10]. Incorporating additional evolutionary steps—including the development of defense and counter-defense mechanisms—will therefore be an interesting extension of our model.

Our analysis provides an intuitive description of pattern formation in our system. At high rates of bacterial or phage proliferation, MIPS is suppressed, and bacteria and phage coexist uniformly across space. By contrast, at sufficiently low phage density and infection rate, bacteria can aggregate via MIPS prior to phage infection, leading to multiple possible outcomes. When phage diffusivity is sufficiently large, stationary finite-sized patterns emerge, with aggregates of bacteria and phage that coexist—in stark contrast to the coarsening observed in conventional MIPS. In this scenario, bacterial death inside aggregates is compensated by a MIPS flux of bacteria from the space between aggregates, where bacteria can proliferate. Conversely, when phage diffusivity is sufficiently small, phage accumulate in and ultimately split or destroy bacterial aggregates, after which continued bacterial proliferation can generate new aggregates—leading to a rich variety of dynamic patterns with intricate spatio-temporal structure. Indeed, the auto-catalytic proliferation mechanism of phage is essential for the emergence of these dynamic patterns; they do not arise in an alternate model in which bacteria are instead killed by a self-secreted toxin.

### Limitations and extensions of our model

Our model provides a foundation for capturing the rich spatio-temporal organization of bacteria-phage populations in nature, and necessarily relies on several simplifications and approximations. For example, we assumed that bacteria undergo instantaneous lysis upon phage infection. However, in reality, bacteria exhibit a latent infection period before lysis, which can influence how phage [9, 75, 76] or bacteria [7] collectively spread. We anticipate that adding this feature to our model would be akin to reducing the phage infection rate and/or burst size, resulting in shifts to phase boundaries but without qualitatively altering the nature of the patterns that emerge.

Another key assumption is that the bacteria aggregate via MIPS. While this canonical model of active matter provides a useful foundation for our work, and MIPS-like behavior has been reported in many different bacterial systems [31– 39], it will not be generally applicable to all bacteria. Additional complexities may arise due to e.g., hydrodynamic, steric, and other mechanical interactions between cells [77– 80], quorum sensing [81, 82], chemotaxis [65, 83, 84], and nutrient-dependent aggregation and proliferation [31, 85]. This study focused on lytic or virulent phages, which infect bacterial cells by injecting genetic material, proliferating inside the cytoplasm, and causing cell lysis to release more phage. However, many natural phages are lysogenic or temperate, meaning that they can integrate their genome into a bacterial cell and thus its progeny *without* lysis. This process renders the cells immune to future infections by the same phage type, but also means they can potentially re-enter the lytic pathway in some future generation. Recent studies indicate that lysogeny can have a marked influence on bacteria-phage dynamics in well-mixed conditions [86]; thus, studying the impact of lysogeny on spatial organization in our model will be an interesting extension of our work.

Previous studies of active mixtures and living systems have demonstrated that noise can impact pattern formation [31, 87– 89]. The presence of noise is inherent to almost all natural environments that bacteria and phage inhabit. Although noise is present in the initial conditions of our simulations, we do not include any additional sources of noise over time. Exploring how the results presented here are modified by the addition of different kinds of noise will be another useful direction for future work.

### Similarities and differences to other nonequilibrium systems

Pattern formation is ubiquitous in nature, arising in both non-living [87, 90–98] and living systems [87, 93, 94, 96, 97, 99–103]. Indeed, systems as diverse as proteins, amoeba, and even animal skin exhibit patterns that are remarkably similar to those revealed by our work [36–38, 90, 91, 95, 102, 104– 117], but which arise from a Turing-like instability due to differential diffusivity between two different species [61]. Our work provides a counterpoint to this paradigm: in our case, patterns do not rely on the Turing mechanism, but instead arise from a combination of aggregation via MIPS and non-reciprocal predator-prey interactions. Moreover, we find that such patterns, once established, persist beyond the phase boundaries predicted by linear stability theory, implying hysteresis. In the same regions, the system exhibits excitability in response to finite perturbations. While many biological systems are excitable via a Turing-type mechanism [104– 110, 118–120], excitability in the context of the non-Turing-type context of bacteria and phage is essentially unexplored. These two phenomena—hysteresis and excitability—emerge as consequences of nonlinear interactions in bacteria-phage populations. Further exploration of the nonlinear phases, phase boundaries, and excitable behavior will therefore be useful future research directions building on our work.

To our knowledge, ours is the first study of the influence of non-reciprocal predator-prey interactions on MIPS. Other forms of non-reciprocity, typically introduced by means of asymmetric cross-diffusive fluxes, have recently been explored in the context of active matter [40–52, 64, 121]. Here, we found a broader range of static and dynamic patterns. Thus, motivated by the comparison between this body of work and ours, unraveling the similarities and differences between the rich patterns and phase behaviors that emerge from different forms of non-reciprocity will be intriguing in future work.

### Implications for biology

As noted above, recent experiments involving swimming and growing bacteria in the presence of lytic phage in a 2D environment have revealed rich spatio-temporal patterns that bear similarities to those revealed by our model [10]. In the experiments, the patterns were characterized by prolonged coexistence of bacteria and phage, which underwent multiple evolutionary cycles. Given that our model does not account for bacteria and phage mutations, it is likely relevant only to short-term patterning, which could then influence longer-term co-evolution. Nevertheless, in both the experiments and in our model, when bacterial collective motility exceeds phage diffusivity, aggregates of bacteria migrate and are then annihilated by phage; the bacteria then require a “refractory” period to regrow, aggregate, and collectively migrate once again. This similarity between our model predictions and experimental observations suggest the tantalizing idea that some of the patterns observed in experiments might be triggered by a coupling between a positive feedback loop that promotes bacterial aggregation—such as MIPS, chemotaxis, or quorum sensing—and a negative feedback loop induced by the inherently non-reciprocal nature of bacteria-phage interactions. Ultimately, incorporating mutations into our model and obtaining the spatial patterns of co-evolved bacteria and phage could provide insights into bacteria-phage co-evolutionary experiments in extended spatial environments.

Our results are not restricted to the case of bacteria and phage; they are also relevant to a broader range of active and living systems that are similarly characterized by non-reciprocal predator-prey interactions. For example, many ecological niches are inhabited by predatory bacteria that prey and feed on other prokaryotic organisms. This is the case of *Myxococcus xanthus*, which preys on *Escherichia coli* by secreting bacteriocins [122], or *Bdellovibrio bacteriovorus*, which consumes Gram-negative bacteria by using pili to penetrate into the periplasm, feed and proliferate in the periplasmic space, and lyse their prey to release offspring [123]. Our results could inform and motivate new experiments on such predator-prey systems in extended spatial environments. On a larger scale, multicellular organisms like fire ants are known to aggregate via a MIPS-like mechanism in laboratory settings [53], yet the effect of predator-prey interactions on these aggregation patterns remains unexplored. By deepening our understanding of how the interactions between an aggregating prey and its predators manifest in intricate spatio-temporal patterns, we hope to inspire and inform experiments in areas as diverse as ecology, phage-based therapies, and proliferating active matter.

## Acknowledgements

A.M.-C. acknowledges support from the Princeton Center for Theoretical Science and the Human Frontier Science Program through the grant LT000035/2021-C. S.S.D. acknowledges support from NSF Grants CBET-1941716, DMR-2011750, and EF-2124863 as well as the Camille Dreyfus Teacher-Scholar and Pew Biomedical Scholars Programs. N.S.W. acknowledges support from the NSF through the Center for the Physics of Biological Function PHY-1734030 and the NIH through grants R01 GM140032 and R01 GM082938. We also thank the Princeton Biomolecular Condensate Program for funding support. We thank Yigal Meir, Carolina Trenado-Yuste, and Hongbo Zhao for thoughtful discussions.

## SUPPLEMENTARY INFORMATION

### Estimation of phage infection rate γ

To estimate the rate at which bacteria lyse due to phage infectivity per phage density *γ*, we first solve the steady-state phage diffusion equation, i.e., *D*_*p*_ **∇**^2^*p* = 0, in spherical coordinates, assuming that a bacterium is a perfectly absorbing sphere of radius *R*_b_. The steady-state phage concentration reads, *p* = *p*_*∞*_(1 −*R*_b_*/r*), where *p*_*∞*_ is a far-field phage concentration. The flux of phage at the sphere’s surface reads 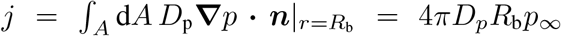 where *A* is the sphere’s surface area. Assuming that a single phage is sufficient to infect and kill a bacterium upon encounter, the bacterium average radius is *R*_b_ ≃ 3*μ*m, and using the estimate of phage diffusivity in Table S1, we obtain an infection rate of *γ* ≃ 9 *×*10^−15^ mL phage^−1^ min^−1^, in good agreement with previous calculations [124, 125].

### Dimensionless equations

Employing the characteristic scales given in the Main Text, the dimensionless versions of Eqs. (1a) and (1b) read:

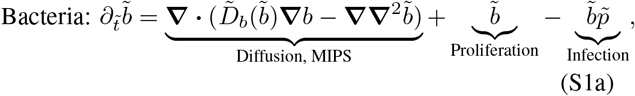

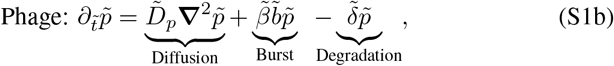

where 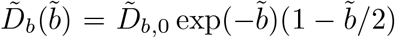 is the dimensionless bacterial effective diffusivity.

### Dimensional stability conditions for the bacteria-phage model

The stability conditions for the linearly-unstable regimes in dimensional variables read:

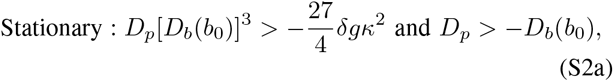

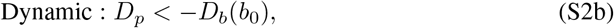

where *D*_*b*_(*b*_0_) = *D*_*b*,0_ exp[−*δα/*(*βγ*)][1− *δα/*(2*βγ*)] is the dimensional effective bacterial diffusivity evaluated at the uniform solution.

### Linear stability analysis in limiting cases

Here we obtain the growth rate of small-amplitude perturbations by taking different limits of the more general dispersion relation, Eq. (7).

#### Slow bacterial and phage diffusion

In the limit of slow bacteria and phage diffusion, 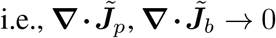, the system reduces to the Lotka-Volterra model [56, 57]. Small perturbations of bacterial and phage densities do not grow over time but rather oscillate with a fixed amplitude that depends on the initial perturbation, which is usually referred to as *stable center* in dynamical systems, and implies a purely imaginary growth rate of perturbations, 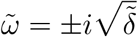, or equivalently 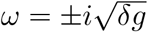

#### Fast phage diffusion and kinetics

In the limit in which phage rapidly relax to a stationary state where diffusion and kinetics are in balance, 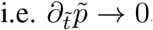, we obtain the following growth rate of perturbations: 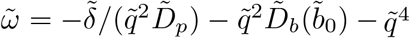 As expected from Fig. **2**a, the growth rate is purely real, and long wavelengths are suppressed which implies arrested coarsening. In the limit 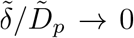, we recover the conventional MIPS growth rate of perturbations, in which perturbations are unstable when 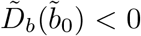, without oscillations and with coarsening over time.

#### Slow phage diffusion

The limit of slow phage diffusion implies 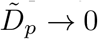. Taking this limit in Eq. (7) yields the following perturbation growth rate:

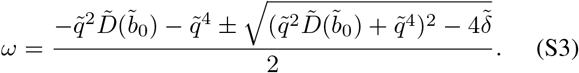

In this limit, the condition for the system to become unstable is the same as for conventional MIPS, 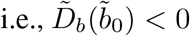. Additionally, long-wavelength modes are always unstable. In this case, certain wavenumbers are also oscillatory as shown by Fig. **2**a in the Main Text, since 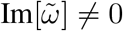. In particular, some long wavelengths are unstable and oscillatory, 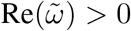 and 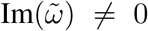. Depending on the parameter values, the wavenumbers that satisfy these two conditions change. In particular:

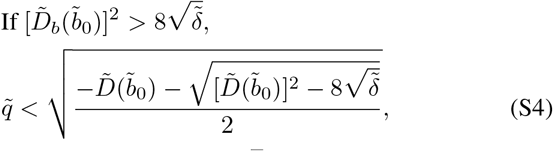

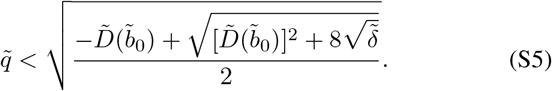

### Bacteria that secrete a self-toxin

Some bacteria are known to secrete self-destructive toxins [72, 73], and more generally bacteria may toxify their environment with waste products [74]. To unravel if the stationary and dynamic patterns described in the Main Text are specific to bacteria-phage non-reciprocal interactions, we investigate a similar model, without phage, but in which bacteria secrete a toxin at rate *ν* (see Fig. **S1**a). The only difference with respect to Eqs. (1) of the Main Text lies in the term describing phage proliferation, which is replaced by a toxin production term proportional to the local density of bacteria only. Thus, the kinetic equations for bacteria and toxin read:

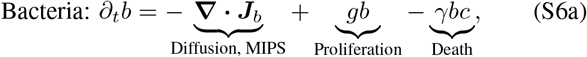

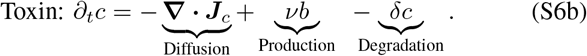

**Table S1.** Estimates of the parameter values.

| Physical parameters | Definition | Range of values |
| --- | --- | --- |
| $g$ | Bacteria proliferation rate | $0.7 \text{ h}^{-1}$ |
| $\alpha$ | Bacteria speed vs. decay rate | $10^{-11} - 10^{-12} \text{ mL cells}^{-1}$ |
| $D_b$ | Bacteria diffusivity | $10^2 - 10^3 \mu\text{m}^2 \text{ s}^{-1}$ |
| $D_p$ | Phage diffusivity | $4 \mu\text{m}^2 \text{ s}^{-1}$ [9] |
| $\delta$ | Phage decay rate | $1.5 \times 10^{-3} - 2 \times 10^{-2} \text{ h}^{-1}$ [126] |
| $\gamma$ | Infection rate | $10^{-14} \text{ mL phage}^{-1} \text{ min}^{-1}$ [126] |
| $\beta$ | Phage burst size | 50-400 [126] |
| Dimensionless parameters |  |  |
| $\tilde{\beta} \equiv \beta\gamma/(\alpha g)$ | Phage proliferation/Bacterial proliferation | 3.9 – 310 |
| $\tilde{\delta} \equiv \delta/g$ | Phage decay rate/Bacterial proliferation | $2 \times 10^{-3} - 3 \times 10^{-2}$ |
| $\tilde{D}_{b,0}/\tilde{D}_p \equiv D_{b,0}/D_p$ | Bacterial diffusivity/Phage diffusivity | 25 – 250 |

In this case, the uniform solution is given by (*b*_0_, *c*_0_) = (*δg/*(*νγ*), *g/γ*). Before carrying out a stability analysis, we reduce the number of parameters via non-dimensionalization. To this end, we choose *t*_c_ = *g*^−1^, *ℓ*_c_ = (*κ/g*)^1*/*4^, *b*_c_ = *α*^−1^, and *c*_c_ = *g/γ* as characteristic time, length, bacterial density, and toxin density scales, respectively. The resulting dimensionless equations are:

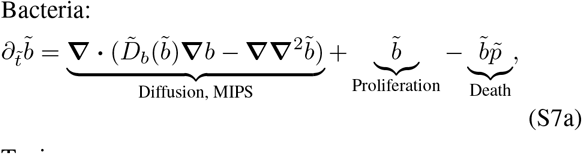

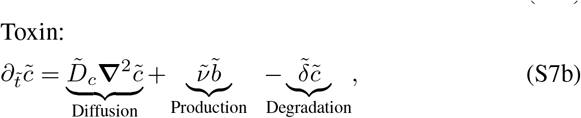

whose uniform solution is 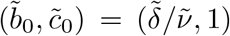. The dimensionless parameters in Eqs. (S7) are:

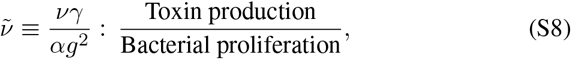

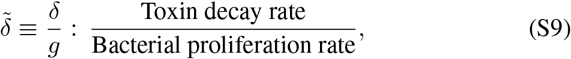

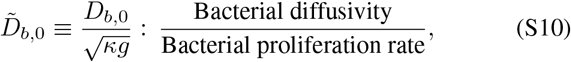

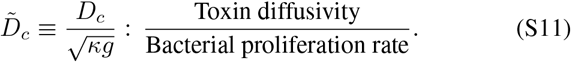

We carry out the same linear stability analysis as performed in the Main Text for the bacteria-phage system. In this case, the dispersion relation reads:

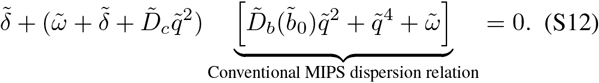

We first obtain the condition under which the system becomes unstable, i.e., 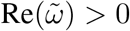for some modes:

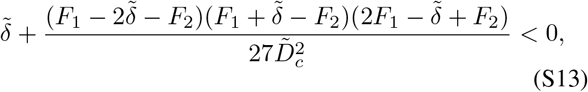

where 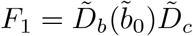and 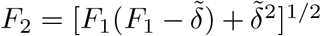. Furthermore, we find that all unstable modes are non-oscillatory, 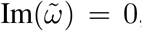, and long wavelengths are stable for all parameter values, as 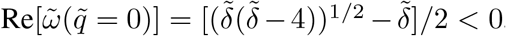, implying that coarsening is arrested in the linear regime.

### Numerical method

To unveil the full nonlinear behavior of Eqs. (1), we perform two-dimensional numerical simulations using the finite-element method. Thus, we write Eqs. (1) in weak form by means of integral scalar products using test functions 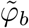 and 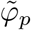 for the bacterial density 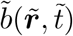 and phage density 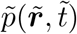 fields, respectively. Using Green identities, we obtain an integral bilinear system of equations for the variables and their test functions:

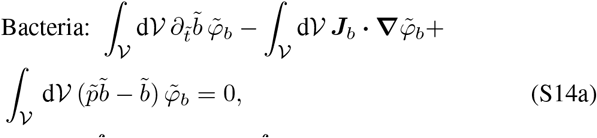

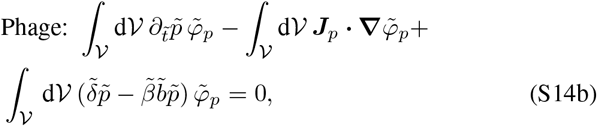

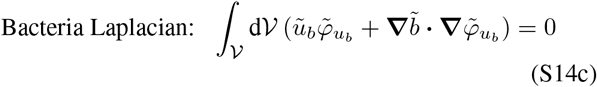

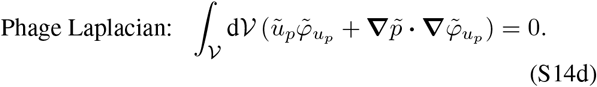

Here𝒱 is the 2D𝒱 domain and d are the surface elements. We impose periodic boundary conditions for both bacteria 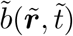 and phage 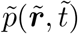. To ensure numerical stability, the equations are discretized in space using second-order Lagrange polynomials and triangular elements for the fields, and evolved in time through a 4th-order variable-step backward differentiation formula method. The relative tolerance of the nonlinear method is always set below 10^−6^.

### Characteristic length scale of the patterns

We obtain the characteristic length scale of the emergent patterns over time from the structure factor as [127–129]:

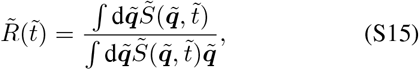

where 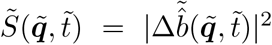 is the structure factor of bacterial amplitude profile 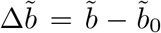, and 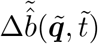Fourier transform.

**Fig. S1.**
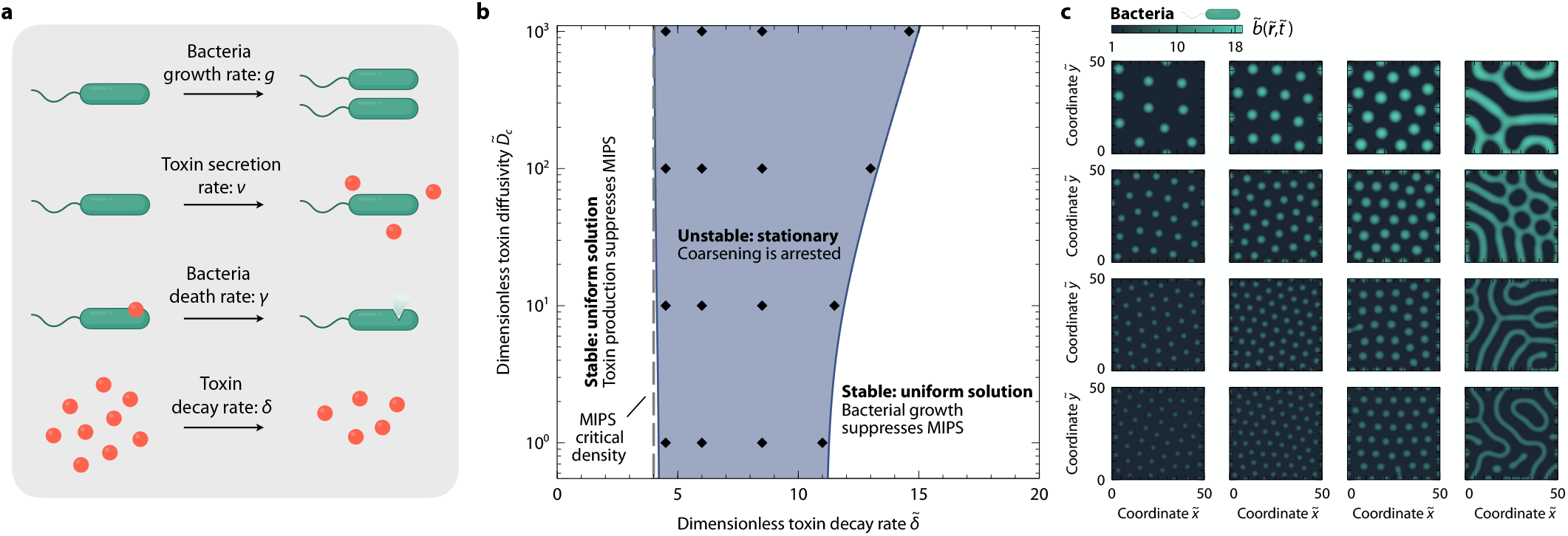
Bacteria-toxin interactions give rise to stationary patterns. **a**. Sketch of the main elements of the model. Bacteria proliferate at a rate *g*. Bacteria secrete a toxin at a rate *ν* that kills bacteria at a rate *γ*. Toxin decays at a rate *δ*. **b**. Linear stability phase diagram as a function of the dimensionless toxin diffusivity 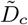 and toxin decay rate 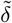, for dimensionless toxin-production-to-bacteria-proliferation ratio and 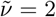 bacterial diffusivity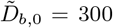. **c**. Two-dimensional plots of and the bacterial density 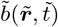 at steady state for different regions of the linear stability phase diagram of panel b as indicated with symbols.

### Characterization of spatio-temporal chaos

Well inside critical boundaries of the region of dynamic instability sh in Fig. **2**a, the patterns that emerge are highly disord and irregular in space and time (Fig. **2**c and e, and Fig. This observation hints that the system could potentially hibit chaotic dynamics. To test this hypothesis, we com the maximum Lyapunov exponent *λ* of the bacterial de fluctuations at different locations in space for the simul shown in Fig. **2**e over a duration of 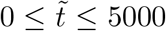 this quantity provides a measure of the system’s sensitivity to s perturbations, with a positive value implying chaotic be ior. Indeed, as shown in Fig. **S2**a, we find that the Lyap exponent is positive in all spatial locations. As an additional characterization of these chaotic dynamics, we follow previous work [130] and compute the temporal correlation function of the bacterial density fluctuations, also for the simulation shown in Fig. **2**e:

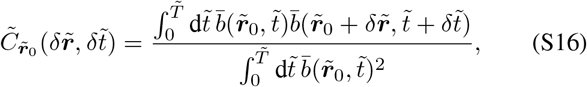

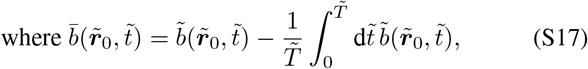

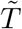 is the temporal window over which 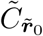 is computed, and 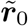 is an arbitrary point in space at which the correlation function is centered. The azimuthally-averaged correlation function 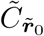 is shown in Fig. **S2**b, which shows that density correlations die away rapidly.

**Fig. S2.**
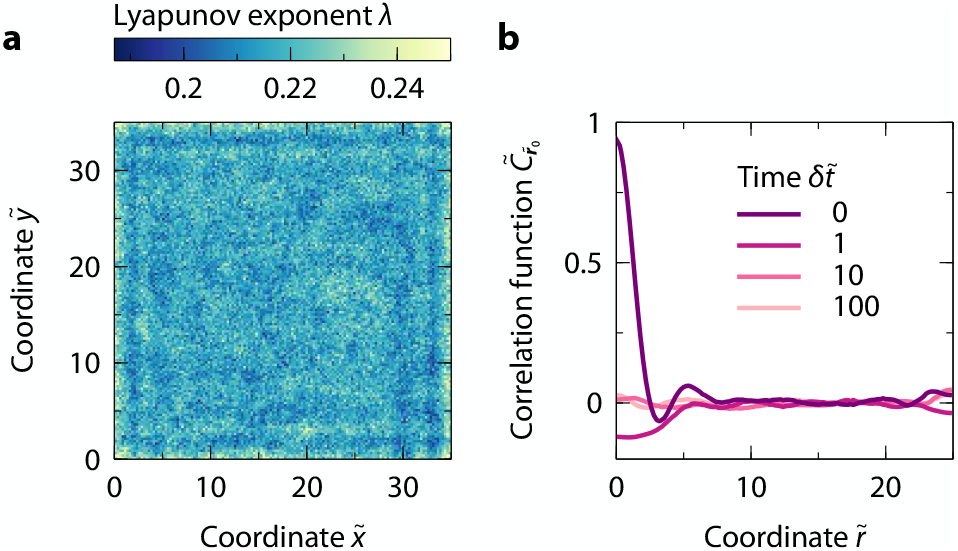
Bacteria-phage dynamic patterns are chaotic. **a**, Maximum Lyapunov exponent *λ* of the bacterial density fluctuations at different locations in the 2D space for the simulation shown in Fig. **2**e. **b**, Azimuthally-averaged correlation function 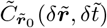 of the bacterial density fluctuations as a function of the radial coordinate 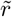 at different time differences 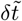, computed at 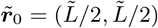 and over a time window of 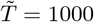.

### Parameter values of Supplementary Movies

The physically reasonable parameter values used in the simulations corresponding to Movies 1-10 are 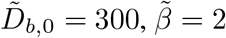 and:

- Movies 1 and 2: 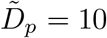, 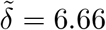.
- Movies 3 and 4: 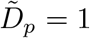, 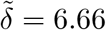.
- Movies 5 and 6: 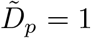, 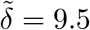.
- Movies 7 and 8: 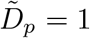, 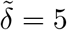.
- Movies 9 and 10: 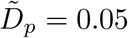, 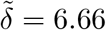.
- Movies 11 and 12: 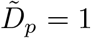, 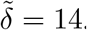.
- Movies 13 and 14: 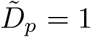, 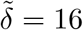.

In all simulations the dimensionless size of the 2D domain is set to 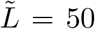, except otherwise specified in the filename. In the simulations corresponding to Movies 11-14, where we show that the system exhibits excitability, instead of initially perturbing the uniform solution with small-amplitude white noise, we introduce a Gaussian perturbation at the center of the 2D spatial domain with amplitude 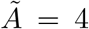 and variance 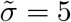:

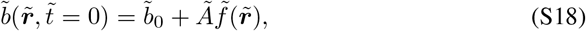

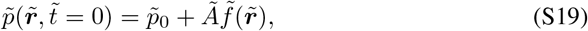

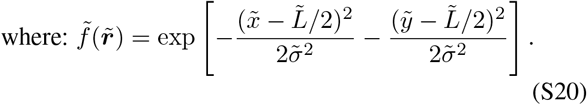

The Supplementary Movies can be found at the following Zenodo repository: https://doi.org/10.5281/zenodo. 8361388

**Fig. S3.**
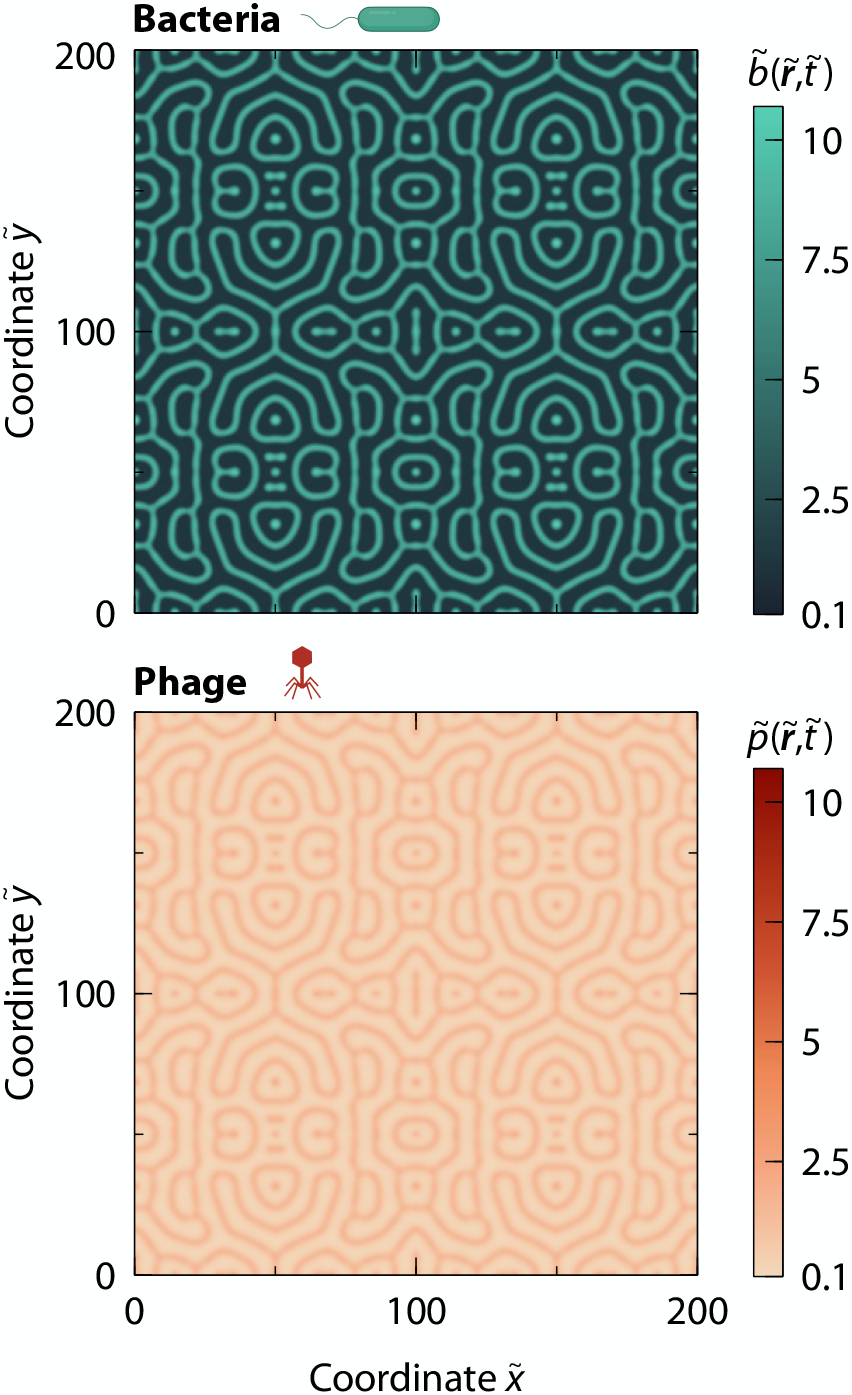
MIPS and non-reciprocal predator-prey relations can give rise to complex stationary patterns. Two-dimensional plot of the bacterial and phage densities at steady state, for a linearly-unstable non-oscillatory case close to the critical boundary in Fig. **2**a, where 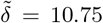 and 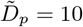. Bacteria and phage coexist in a network of interconnected stripes.

## References

[1] H. G. Hampton, B. N. J. Watson, and P. C. Fineran, Nature 577, 327 (2020).

[2] A. Chevallereau, B. J. Pons, S. van Houte, and E. R. Westra, Nat. Rev. Microbiol. 20, 49 (2022).

[3] M. LeRoux and M. T. Laub, Annu. Rev. Microbiol. 76, 21 (2022).

[4] M. Lourenço, L. Chaffringeon, Q. Lamy-Besnier, T. Pédron, P. Campagne, C. Eberl, M. Bérard, B. Stecher, L. Debarbieux, and L. De Sordi, Cell Host Microbe 28, 390 (2020).

[5] B. Koskella, C. A. Hernandez, and R. M. Wheatley, Annu. Rev. Virol. 9, 57 (2022).

[6] M. Breitbart, C. Bonnain, K. Malki, and N. Sawaya, Nat. Microbiol. 3, 754 (2018).

[7] D. Ping, T. Wang, D. T. Fraebel, S. Maslov, K. Sneppen, and S. Kuehn, ISME J. 14, 2007 (2020).

[8] E. L. Simmons, M. C. Bond, B. Koskella, K. Drescher, V. Bucci, and C. D. Nadell, mSystems 5, e00877 (2020).

[9] M. Hunter, N. Krishnan, T. Liu, W. Möbius, and D. Fusco, Phys. Rev. X 11, 021066 (2021).

[10] E. Shaer Tamar and R. Kishony, Nat. Commun. 13, 7971 (2022).

[11] J. B. Winans, B. R. Wucher, and C. D. Nadell, PLoS Biol. 20, e3001913 (2022).

[12] R. S. Eriksen, F. A. N. Larsen, S. L. Svenningsen, K. Sneppen, and N. Mitarai, bioRxiv, 2023 (2023).

[13] P. H. Thrall and J. J. Burdon, Plant Pathol. 51, 169 (2002).

[14] F. Débarre, S. Lion, M. Van Baalen, and S. Gandon, Am. Nat. 179, 52 (2012).

[15] J. O. Haerter and K. Sneppen, MBio 3, 10 (2012).

[16] E. L. Simmons, K. Drescher, C. D. Nadell, and V. Bucci, ISME J. 12, 531 (2018).

[17] R. S. Eriksen, N. Mitarai, and K. Sneppen, Sci. Rep. 10, 3154 (2020).

[18] H. I. Egilmez, A. Y. Morozov, and E. E. Galyov, Sci. Rep. 11, 4382 (2021).

[19] B. Kerr, C. Neuhauser, B. J. M. Bohannan, and A. M. Dean, Nature 442, 75 (2006).

[20] L. Chao, B. R. Levin, and F. M. Stewart, Ecology 58, 369 (1977).

[21] R. E. Lenski and B. R. Levin, Am. Nat. 125, 585 (1985).

[22] B. R. Levin and J. J. Bull, Am. Nat. 147, 881 (1996).

[23] Y. Wei, A. Kirby, and B. R. Levin, Am. Nat. 178, 715 (2011).

[24] M. F. Marston, F. J. Pierciey Jr, A. Shepard, G. Gearin, J. Qi, C. Yandava, S. C. Schuster, M. R. Henn, and J. B. H. Martiny, Proc. Natl. Acad. Sci. U.S.A. 109, 4544 (2012).

[25] A. Betts, O. Kaltz, and M. E. Hochberg, Proc. Natl. Acad. Sci. U.S.A. 111, 11109 (2014).

[26] S. van Houte, A. K. E. Ekroth, J. M. Broniewski, H. Chabas, B. Ashby, J. Bondy-Denomy, S. Gandon, M. Boots, S. Paterson, A. Buckling, et al., Nature 532, 385 (2016).

[27] >A. Oromí-Bosch, J. Antani, and P. Turner, Annu. Rev. Virol. 10 (2023).

[28] M. J. Schnitzer, Phys. Rev. E 48, 2553 (1993).

[29] J. Tailleur and M. E. Cates, Phys. Rev. Lett. 100, 218103 (2008).

[30] M. E. Cates and J. Tailleur, Annu. Rev. Condens. Matter Phys. 6, 219 (2015).

[31] G. Liu, A. Patch, F. Bahar, D. Yllanes, R. D. Welch, M. C. Marchetti, S. Thutupalli, and J. W. Shaevitz, Phys. Rev. Lett. 122, 248102 (2019).

[32] C. Liu, X. Fu, L. Liu, X. Ren, C. K. L. Chau, S. Li, L. Xiang, H. Zeng, G. Chen, L. H. Tang, et al., Science 334, 238 (2011).

[33] X. Fu, L. H. Tang, C. Liu, J. D. Huang, T. Hwa, and P. Lenz, Phys. Rev. Lett. 108, 198102 (2012).

[34] A. I. Curatolo, N. Zhou, Y. Zhao, C. Liu, A. Daerr, J. Tailleur, and J. Huang, Nat. Phys. 16, 1152 (2020).

[35] I. Grobas, M. Polin, and M. Asally, Elife 10, e62632 (2021).

[36] D. E. Woodward, R. Tyson, M. R. Myerscough, J. D. Murray, E. O. Budrene, and H. C. Berg, Biophys. J. 68, 2181 (1995).

[37] M. P. Brenner, L. S. Levitov, and E. O. Budrene, Biophys. J. 74, 1677 (1998).

[38] M. E. Cates, D. Marenduzzo, I. Pagonabarraga, and J. Tailleur, Proc. Natl. Acad. Sci. U.S.A. 107, 11715 (2010).

[39] S. Park, P. M. Wolanin, E. A. Yuzbashyan, H. Lin, N. C. Darnton, J. B. Stock, P. Silberzan, and R. Austin, Proc. Natl. Acad. Sci. U.S.A. 100, 13910 (2003).

[40] M. Fruchart, R. Hanai, P. Littlewood, and V. Vitelli, Nature 592, 363 (2021).

[41] J. Zhang, R. Alert, J. Yan, N. S. Wingreen, and S. Granick, Nat. Phys. 17, 961 (2021).

[42] S. Shankar, A. Souslov, M. Bowick, M. Marchetti, and V. Vitelli, Nat. Rev. Phys. 4, 380 (2022).

[43] M. Nagy, Z. Ákos, D. Biro, and T. Vicsek, Nature 464, 890 (2010).

[44] D. Yllanes, M. Leoni, and M. C. Marchetti, New J. Phys. 19, 103026 (2017).

[45] L. L. Bonilla and C. Trenado, Phys. Rev. E 99, 012612 (2019).

[46] F. A. Lavergne, H. Wendehenne, T. Bäuerle, and C. Bechinger, Science 364, 70 (2019).

[47] N. Uchida and R. Golestanian, Phys. Rev. Lett. 104, 178103 (2010).

[48] B. C. van Zuiden, J. Paulose, W. T. M. Irvine, D. Bartolo, and Vitelli, Proc. Natl. Acad. Sci. U.S.A. 113, 12919 (2016).

[49] C. Coulais, D. Sounas, and A. Alu, Nature 542, 461 (2017).

[50] M. Brandenbourger, X. Locsin, E. Lerner, and C. Coulais, Nat. Commun. 10, 4608 (2019).

[51] Z. You, A. Baskaran, and M. C. Marchetti, Proc. Natl. Acad. Sci. U.S.A. 117, 19767 (2020).

[52] S. Saha, J. Agudo-Canalejo, and R. Golestanian, Phys. Rev. X 10, 041009 (2020).

[53] C. Anderson and A. Fernandez-Nieves, Nat. Commun. 13, 6710 (2022).

[54] M. Fruchart, C. Scheibner, and V. Vitelli, Annu. Rev. Condens. Matter Phys. 14, 471 (2023).

[55] O. Hallatschek, S. S. Datta, K. Drescher, J. Dunkel, J. Elgeti, B. Waclaw, and N. S. Wingreen, Nat. Rev. Phys. (2023).

[56] A. J. Lotka, Proc. Natl. Acad. Sci. U.S.A. 6, 410 (1920).

[57] V. Volterra, Variazioni e fluttuazioni del numero d’individui in specie animali conviventi, Vol. 2 (Societá anonima tipografica” Leonardo da Vinci”, 1927).

[58] M. E. Cates and J. Tailleur, Europhys. Lett. 101, 20010 (2013).

[59] G. Gonnella, D. Marenduzzo, A. Suma, and A. Tiribocchi, C. R. Phys. 16, 316 (2015).

[60] T. Grafke, M. E. Cates, and E. Vanden-Eijnden, Phys. Rev. Lett. 119, 188003 (2017).

[61] A. M. Turing, Philos. Trans. R. Soc. Lond., B, Biol. Sci. 237, 37 (1952).

[62] P. W. Voorhees, J. Stat. Phys. 38, 231 (1985).

[63] B. Gouveia, Y. Kim, J. W. Shaevitz, S. Petry, H. A. Stone, and C. P. Brangwynne, Nature 609, 255 (2022).

[64] F. Brauns and M. C. Marchetti, arXiv preprint arXiv:2306.08868 (2023).

[65] H. Zhao, A. Košmrlj, and S. S. Datta, Phys. Rev. Lett. 131, 118301 (2023).

[66] J. W. Cahn and J. E. Hilliard, J. Chem. Phys. 28, 258 (1958).

[67] J. W. Cahn, J. Chem. Phys. 30, 1121 (1959).

[68] J. W. Cahn and J. E. Hilliard, J. Chem. Phys. 31, 688 (1959).

[69] J. W. Cahn, J. Chem. Phys. 42, 93 (1965).

[70] J. Berry, C. Brangwynne, and M. Haataja, Rep. Prog. Phys. 81, 046601 (2018).

[71] C. Weber, D. Zwicker, F. Jülicher, and C. Lee, Rep. Prog. Phys. 82, 064601 (2019).

[72] P. F. Popp and T. Mascher, J. Mol. Biol. 431, 4656 (2019).

[73] X. Jin, S. An, W. Kightlinger, J. Zhou, and S. H. Hong, AIChE J. 67, e17466 (2021).

[74] C. Ratzke, J. Denk, and J. Gore, Nat. Ecol. Evol. 2, 867 (2018).

[75] J. Yin and J. McCaskill, Biophys. J. 61, 1540 (1992).

[76] J. Fort and V. Méndez, Phys. Rev. Lett. 89, 178101 (2002).

[77] M. Theers, E. Westphal, K. Qi, R. G. Winkler, and G. Gompper, Soft matter 14, 8590 (2018).

[78] R. M. Navarro and S. M. Fielding, Soft Matter 11, 7525 (2015).

[79] J. N. Wilking, T. E. Angelini, A. Seminara, M. P. Brenner, and D. A. Weitz, MRS Bull. 36, 385 (2011).

[80] M. E. Black and J. W. Shaevitz, Phys. Rev. Lett. 130, 218402 (2023).

[81] J. A. Moore-Ott, S. Chiu, D. B. Amchin, T. Bhattacharjee, and S. S. Datta, eLife 11, e76380 (2022).

[82] W. J. M. Ridgway, M. P. Dalwadi, P. Pearce, and S. J. Chapman, bioRxiv, 2023 (2023).

[83] T. Bhattacharjee, D. B. Amchin, R. Alert, J. A. Ott, and S. S. Datta, eLife 11, e71226 (2022).

[84] F. Fadda, D. A. Matoz-Fernandez, R. van Roij, and S. Jabbari-Farouji, Soft Matter 19, 2297 (2023).

[85] A. Martínez-Calvo, T. Bhattacharjee, R. K. Bay, H. N. Luu, A. M. Hancock, N. S. Wingreen, and S. S. Datta, Proc. Natl. Acad. Sci. U.S.A. 119, e2208019119 (2022).

[86] O. Kimchi, Y. Meir, and N. S. Wingreen, bioRxiv, 2022 (2022).

[87] F. Sagués, J. M. Sancho, and J. García-Ojalvo, Rev. Mod. Phys. 79, 829 (2007).

[88] E. Tjhung, C. Nardini, and M. E. Cates, Phys. Rev. X 8, 031080 (2018).

[89] Y. I. Li and M. E. Cates, Eur. Phys. J. E 44, 119 (2021).

[90] B. P. Belousov, Ref. Radiats. Med. (1958).

[91] A. M. Zhabotinsky, Biofizika 9, 306 (1964).

[92] D. A. Kessler, J. Koplik, and H. Levine, Adv. Phys. 37, 255 (1988).

[93] M. C. Cross and P. C. Hohenberg, Rev. Mod. Phys. 65, 851 (1993).

[94] E. Ben-Jacob, I. Cohen, and H. Levine, Adv. Phys. 49, 395 (2000).

[95] V. K. Vanag and I. R. Epstein, Science 294, 835 (2001).

[96] J. A. Acebrón, L. L. Bonilla, C. J. P. Vicente, F. Ritort, and R. Spigler, Rev. Mod. Phys. 77, 137 (2005).

[97] M. C. Marchetti, J. F. Joanny, S. Ramaswamy, T. B. Liverpool, J. Prost, M. Rao, and R. A. Simha, Rev. Mod. Phys. 85, 1143 (2013).

[98] Y. Fuseya, H. Katsuno, K. Behnia, and A. Kapitulnik, Nat. Phys. 17, 1031 (2021).

[99] J. A. Thomson, “On growth and form,” (1917).

[100] A. J. Koch and H. Meinhardt, Rev. Mod. Phys. 66, 1481 (1994).

[101] E. Ben-Jacob, O. Schochet, A. Tenenbaum, I. Cohen, A. Czirok, and T. Vicsek, Nature 368, 46 (1994).

[102] S. Kondo and T. Miura, Science 329, 1616 (2010).

[103] B. D’Aquino, Bull. Am. Phys. Soc/ (2023).

[104] K. C. Huang, Y. Meir, and N. S. Wingreen, Proc. Natl. Acad. Sci. U.S.A. 100, 12724 (2003).

[105] K. C. Huang and N. S. Wingreen, Phys. Biol. 1, 229 (2004).

[106] M. Loose, E. Fischer-Friedrich, J. Ries, K. Kruse, and P. Schwille, Science 320, 789 (2008).

[107] V. Ivanov and K. Mizuuchi, Proc. Natl. Acad. Sci. U.S.A. 107, 8071 (2010).

[108] M. Loose, E. Fischer-Friedrich, C. Herold, K. Kruse, and P. Schwille, Nat. Struct. Mol. Biol. 18, 577 (2011).

[109] A. G. Vecchiarelli, M. Li, M. Mizuuchi, L. C. Hwang, Y. Seol, K. C. Neuman, and K. Mizuuchi, Proc. Natl. Acad. Sci. U.S.A. 113, E1479 (2016).

[110] B. Ramm, A. Goychuk, A. Khmelinskaia, P. Blumhardt, H. Eto, K. A. Ganzinger, E. Frey, and P. Schwille, Nat. Phys. 17, 850 (2021).

[111] R. Alert, A. Martínez-Calvo, and S. S. Datta, Phys. Rev. Lett. 128, 148101 (2022).

[112] J. T. Bonner, Q. Rev. Biol. 32, 232 (1957).

[113] P. Foerster, S. C. Müller, and B. Hess, Dev. 109, 11 (1990).

[114] K. J. Lee, E. C. Cox, and R. E. Goldstein, Phys. Rev. Lett. 76, 1174 (1996).

[115] J. Lauzeral, J. Halloy, and A. Goldbeter, Proc. Natl. Acad. Sci. U.S.A. 94, 9153 (1997).

[116] M. Falcke and H. Levine, Phys. Rev. Lett. 80, 3875 (1998).

[117] J. T. Bonner, Cellular slime molds, Vol. 2127 (Princeton University Press, 2015).

[118] W. M. Bement, M. Leda, A. M. Moe, A. M. Kita, M. E. Larson, A. E. Golding, C. Pfeuti, K. C. Su, A. L. Miller, A. B. Goryachev, et al., Nat. Cell Biol. 17, 1471 (2015).

[119] T. H. Tan, J. Liu, P. W. Miller, M. Tekant, J. Dunkel, and N. Fakhri, Nat. Phys. 16, 657 (2020).

[120] M. C. Wigbers, T. H. Tan, F. Brauns, J. Liu, S. Z. Swartz, E. Frey, and N. Fakhri, Nat. Phys. 17, 578 (2021).

[121] A. Dinelli, J. O’Byrne, A. Curatolo, Y. Zhao, P. Sollich, and J. Tailleur, arXiv preprint arXiv:2203.07757 (2022).

[122] R. R. Nair, M. Vasse, S. Wielgoss, L. Sun, Y.-T. N. Yu, and G. J. Velicer, Nat. Commun. 10, 4301 (2019).

[123] R. E. Sockett, Annu. Rev. Microbiol. 63, 523 (2009).

[124] A. L. Koch, Biochim. Biophys. Acta 39, 311 (1960).

[125] A. G. Murray and G. A. Jackson, Mar. Ecol. Prog. Ser., 103 (1992).

[126] M. De Paepe and F. Taddei, PLoS Biol. 4, e193 (2006).

[127] H. Furukawa, Phys. Rev. E 61, 1423 (2000).

[128] V. M. Kendon, M. E. Cates, I. Pagonabarraga, J. C. Desplat, and P. Bladon, J. Fluid Mech. 440, 147 (2001).

[129] S. Mao, D. Kuldinow, M. P. Haataja, and A. Košmrlj, Soft Matter 15, 1297 (2019).

[130] R. Ramaswamy and F. Jülicher, Sci. Rep. 6, 20838 (2016).

